# Vegetation and seed bank dynamics highlight the importance of post-restoration management in sown grasslands

**DOI:** 10.1101/2020.01.20.913426

**Authors:** Orsolya Valkó, Balázs Deák, Péter Török, Katalin Tóth, Réka Kiss, András Kelemen, Tamás Miglécz, Judit Sonkoly, Béla Tóthmérész

## Abstract

Sowing grass seeds generally supports the rapid development of a closed perennial vegetation, which makes the method universally suitable for fast and effective landscape-scale restoration of grasslands. However, sustaining the recovered grasslands, and increasing their diversity is a challenging task. Understanding the role of seed bank compositional changes and vegetation dynamics contributes to designating management regimes that support the establishment of target species and suppress weeds. Our aim was to reveal the effect of post-restoration management on the vegetation and seed bank dynamics in grasslands restored in one of the largest European landscape-scale restoration projects. Eight years after restoration we sampled the vegetation and seed bank in a total of 96 plots located in 12 recovered grasslands in the Great Hungarian Plain. In each recovered grassland stand we designated a mown (mown from Year 1 to Year 8) and an abandoned sample site (mown from Year 1 to Year 3 then abandoned from Year 4 to Year 8). Mown and abandoned sites showed divergent vegetation and seed bank development. Abandonment led to the decline of sown grasses and higher cover of weeds, especially in the alkaline grasslands. Our study confirmed that seed bank has a limited contribution to the maintenance of biodiversity in both grassland types. We found that five years of abandonment had a larger effect on the seed bank than on the vegetation. We stress that long-term management is crucial for controlling the emergence of the weeds from their dense seed bank in restored grasslands.

**Implications for practice:**

- Seed sowing of grass mixtures can be a feasible tool for restoring grasslands at large scales. However, the developed vegetation usually has low biodiversity and a high seed density of weeds is typical in the soil seed bank even several years after the restoration. Therefore, post-restoration management is necessary for suppressing weeds both aboveground and belowground.
- We recommend to design the long-term management of the sites subjected to grassland restoration already in the planning phase of the restoration projects and ensure that the management plan is ecologically and economically feasible.
- We recommend to complement the monitoring of vegetation with the analysis of soil seed bank for evaluating restoration success.

## Introduction

The restoration of degraded ecosystems is an important strategy to mitigate the negative impacts of human activities on Earth. Grassland restoration is widely applied in nature conservation to increase landscape connectivity, create habitats for plants and animals and restore important ecosystem functions and services (Cole et al. 2019). The best practices for the fast and successful restoration of grasslands characterised by high cover of perennial grasses and low cover of weeds are well developed and widely applied (Kiehl et al. 2010; Török et al. 2011).

In many cases, introduction of seeds in restoration sites is crucial for guaranteeing restoration success, and to ensure the colonisation of the target grassland species. Seed sowing is especially recommended in large restoration sites in human-modified landscapes and in areas subjected to long-lasting or severe degradation, where the restoration potential of seed bank is limited (Török et al. 2018). Dry grasslands in general have a low-density seed bank, characterised by transient seeds, and containing just a few persistent seeds of only a few typical grassland species (Bossuyt & Honnay 2008; Kiss et al. 2016). Therefore, seed sowing is a widely applied propagule introduction method in dry grassland restoration projects. However, the availability of seed material from regional provenance is often a major limiting factor in restoration (de Vitis et al. 2017); thus, especially in large-scale projects only a limited number of target species can be introduced (Valkó et al. 2016a).

Sowing a grass-dominated seed mixture guarantees a directed vegetation development and a cost-effective way of restoration (Kiehl et al. 2010; Török et al. 2011; Valkó et al. 2016b). In such projects the most challenging task is to select the proper species from local provenance and proper density (van der Mijnsbrugge et al. 2010; de Vitis et al. 2017). After sowing the proper seed mixture, we can expect a fast and successful grassland recovery (Baer et al. 2002; Deal et al. 2014; Török et al. 2010). Even though the initial vegetation after sowing is usually characterised by weeds emerging from the seed bank of the formerly degraded areas, sown grasses can competitively exclude them from the aboveground vegetation after two or three years (Török et al. 2010). Therefore, if the seed material and proper machinery is provided, seed sowing can be a universally feasible tool for restoring basic grassland vegetation even on large spatial scales.

The long-term maintenance of restored grasslands is a more challenging task. First, low-diversity communities in general are more sensitive to disturbances because they are less stable than high-diversity communities (Oliver et al. 2015). Greater species richness promotes stability, because there are high number of species that respond differently to the environmental fluctuations, so the decline of one of them could be compensated by the strengthening of another one (Lepš 2004). Second, the legacy of the former degradation, especially in the form of the seed bank of weeds, acts as a threat for future degradation in the species-poor restored grasslands (Halassy 2001; Walker et al. 2004; Török et al. 2012). Finally, the dense grass sward hampers the establishment of target grassland species, but if the grassland is severely disturbed, there is a higher chance for the establishment of the weeds due to their increased propagule availability (Valkó et al. 2016a; Klaus et al. 2018). Considering these threats, it is crucial to develop long-term management strategies to mitigate the degradation of the restored grasslands (Kelemen et al. 2014). Regular mowing or grazing is essential to control weed encroachment and litter accumulation and also for creating establishment microsites for target species (Tälle et al. 2016).

In this study we tested the effects of post-restoration management (mowing vs. abandonment) on the vegetation and seed bank of alkaline and loess grasslands, which were restored during one of the largest grassland restoration projects in Europe. Alkaline grasslands are typical on nutrient-poor, saline soils, and usually have a species-poor vegetation (Deák et al. 2014), but a diverse and dense seed bank (Valkó et al. 2014). Loess grasslands are typical on fertile chernozem soils and are characterised by high plant diversity in the aboveground vegetation (Kelemen et al. 2013), and low seed density and diversity in the seed bank (Tóth & Hüse 2014). This study system offers a unique opportunity for testing the effects of post-restoration management on the vegetation and seed bank of restored grasslands and the dependence of these effects on the grassland type.

We tested the following hypotheses: (i) Abandoned restored grasslands are characterised by lower species richness, lower cover of sown perennial grasses, and higher cover of weeds compared to mown ones (Kelemen et al. 2014). (ii) The effects of abandonment depend on the grassland type and we expect that, due to their low density and low diversity seed bank, restored loess grasslands are more sensitive to abandonment than restored alkaline grasslands (Tóth & Hüse 2014; Valkó et al. 2014). (iii) The effect of abandonment is more pronounced in the vegetation of the restored grasslands than in the seed bank as vegetation dynamics are generally faster than seed bank dynamics (Miao et al. 2016).

## Materials and Methods

### Study area

Our study area is in the Hortobágy National Park (Great Hungarian Plain), near Tiszafüred and Egyek towns. The climate of the region is moderately continental with a mean annual precipitation of 550 mm and a mean temperature of 9.5 °C (Lukács et al. 2015). The National Park holds one of the largest remaining natural open landscapes in Europe, characterised by a diverse mosaic of loess and alkaline grasslands, meadows and wetlands (Deák et al. 2014). Because of their good-quality chernozemic soils, many stands of loess grasslands have been converted to arable fields, and in some regions, large stands of alkaline grasslands with less fertile meadow solonetz soil were also ploughed. The most typical crop plants in the region are alfalfa (*Medicago sativa*), sunflower (*Helianthus annuus*) and wheat (*Triticum aestivum*) (Török et al. 2012).

### Restoration project

In the study area, in total 760 hectares of grasslands were restored on former croplands, which was one of the largest grassland restoration projects in Europe (LIFE 04 NAT HU 119). The aim of the landscape-scale restoration project was to create buffer zones around wetlands and to restore the historical landscape connectivity (Lengyel et al. 2012). Two types of grass seed mixtures were sown after soil preparation in a density of 25kg/ha in October 2005 (Deák et al. 2011). To restore alkaline grasslands, on lower-elevated (<90 m a.s.l.) sites an ‘alkaline seed mixture’ containing the seeds of *Festuca pseudovina* (66%) and *Poa angustifolia* (34%) was sown. To restore loess grasslands, on higher-elevated (>90 m a.s.l.) sites the sown ‘loess seed mixture’ contained the seeds of *Festuca rupicola* (40%), *Poa angustifolia* (30%) and *Bromus inermis* (30%) (Török et al. 2010). These perennial grass species were selected because they are typical in the region, their seeds were available from regional provenance, and they are good competitors which can suppress weeds.

### Vegetation sampling

We selected twelve restored grasslands for the study, seven of them were alkaline and five were loess grasslands. We designated two 5 m × 5 m-sized sites in each grassland. One of the sites was ‘mown’ and has been mown every year by hand in the middle of June, from Year 1 to Year 8. The other site was ‘abandoned’ and was only mown between Year 1 and Year 3, but mowing was stopped from Year 4 onwards. This way we tested the effects of the abandonment of post-restoration management on the vegetation and seed banks of restored grasslands. In each site we designated four 1 m × 1 m-sized permanent plots, where we recorded vegetation data and sampled soil seed banks. For this study, we used data from Year 8. During the vegetation survey, the percentage cover scores of vascular plants were recorded in early June in the 1 m × 1 m plots.

### Seed bank sampling

Soil seed bank was sampled in each plot in Year 8 in late March, after snowmelt. Three soil cores (4 cm diameter, 10 cm depth) were drilled per plot. The three cores originated from the same plot were pooled and processed together. We concentrated seed bank samples according to the protocol of ter Heerdt et al. (1996) for washing out fine mineral and organic particles and reducing sample volume. Rough plant particles were retained on a coarse mesh (2.8 mm), while seeds were retained using a fine mesh (0.2 mm). Concentrated samples were spread in a thin layer on the surface of steam-sterilised potting soil in germination boxes. Samples were germinated under natural light conditions in a greenhouse from early April until early November. Samples were watered regularly, but from mid-July to mid-August we included a drought period when we did not water the pots in order to break dormancy of ungerminated seeds. Germinated seedlings were regularly counted and identified, while unidentifiable seedlings were transplanted and grown until they developed diagnostic features. Accidental air-borne seed contamination was monitored in sample-free control trays filled with steam-sterilized potting soil.

### Data processing

We pooled *Typha latifolia* and *T. angustifolia* as *Typha angustifolia* (in total 193 seedlings) in the analyses as we could not distinguish the germinated specimens due to the lack of flowering. We considered adventive species (e.g. *Conyza canadensis*), ruderal competitors (e.g. *Cirsium arvense*) and weed species (e.g. *Descurainia sophia*) as weeds based on the social behaviour type classification system of Borhidi (1995). We considered unsown generalist, competitor and specialist species typical to grassland habitats as unsown target species (Borhidi 1995). Sown grasses were analysed separately from naturally established unsown target species. We calculated the Jaccard similarity of the species composition of the vegetation and seed bank for each plot.

Effects of ‘management’ (mown, abandoned), ‘grassland type’ (alkaline, loess) and the interaction of ‘management’ and ‘grassland type’ on the vegetation and seed bank characteristics were analysed by Generalized Linear Models in SPSS 20.0 (Zuur et al. 2009). Site was included as random factor. Dependent variables for the vegetation were total species richness, and the cover of annual species, perennial species, weeds, sown grasses and unsown target species. For the seed bank, we included total species richness, and the seedling number of annual species, perennial species, weeds, sown grasses and unsown target species. We identified indicator species of the vegetation and seed bank with the IndVal procedure (Dufrêne & Legendre 1997), using the ‘labdsv’ package in an R environment (R Core Team 2016). Species composition of the vegetation and seed bank was visualised by DCA ordination (detrended correspondence analysis), based on relative abundance data of species using CANOCO 5 (Ter Braak & Šmilauer 2012).

## Results

### Vegetation characteristics

Total species richness was not affected by management and grassland type (Table 1, Figure 1a). The cover of unsown target species was similarly low regardless of management and grassland type (Table 1, Figure 1b). The cover of weeds was affected by the management and the grassland type (Table 1). The highest cover of weeds was recorded in the alkaline grasslands and in the abandoned sites (Figure 1c). The cover of sown grasses was affected by management and the interaction of management and grassland type (Table 1); the highest values were detected in the mown sites and the lowest cover of sown grasses was detected in the abandoned alkaline grasslands (Figure 1d).

**Table 1.**
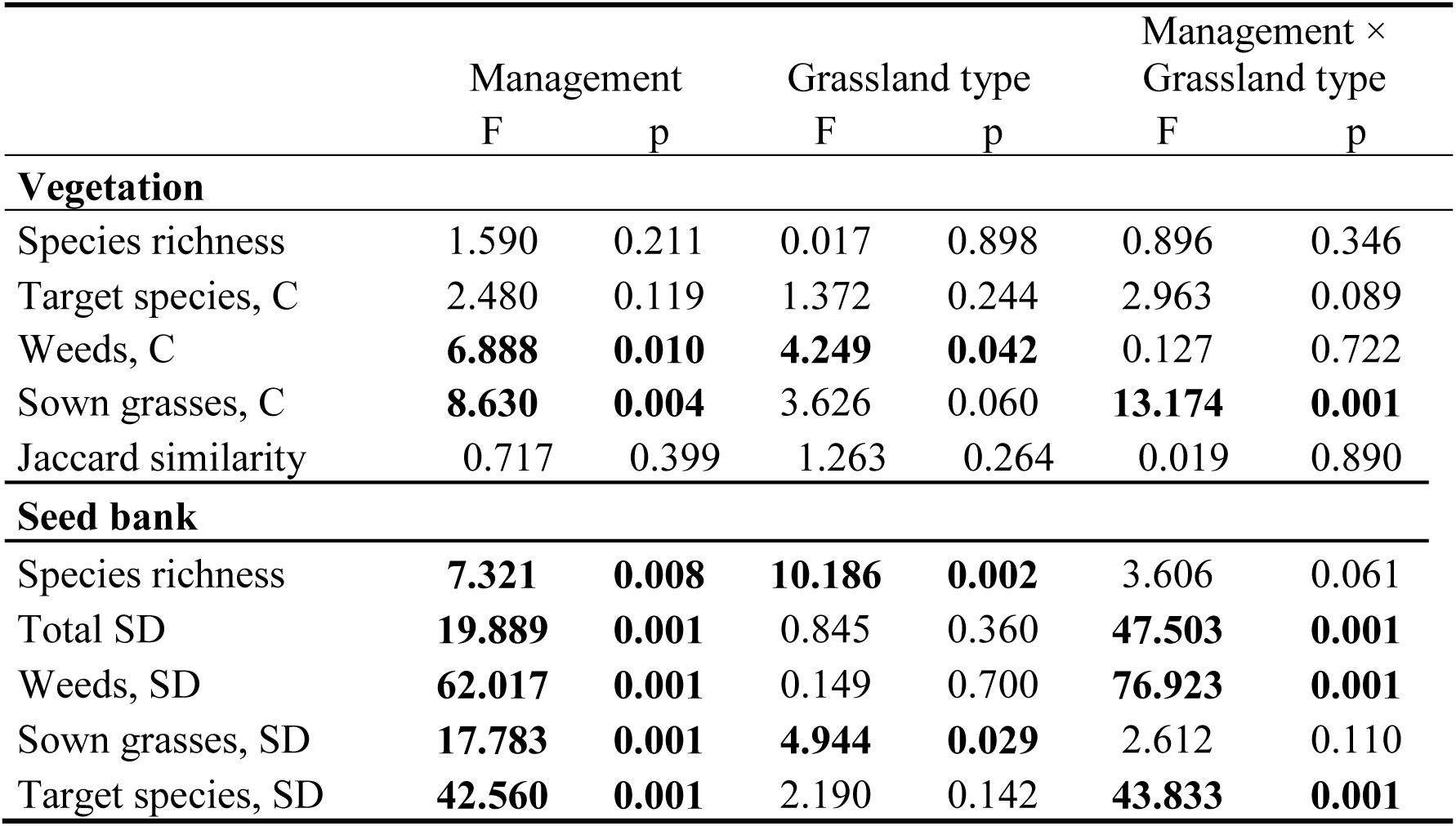
The effect of management (mown/abandoned), grassland type (alkaline/loess) and their interaction on the vegetation and seed bank characteristics of the restored grasslands (Generalized Linear Mixed Models). Notations: C – cover; SD – seed density. Jaccard similarity was calculated between the species composition of the vegetation and the seed bank. Significant differences are marked with boldface.

|  | Management |  | Grassland type |  | Management ×<br>Grassland type |  |
| --- | --- | --- | --- | --- | --- | --- |
|  | F | p | F | p | F | p |
| **Vegetation** |  |  |  |  |  |  |
| Species richness | 1.590 | 0.211 | 0.017 | 0.898 | 0.896 | 0.346 |
| Target species, C | 2.480 | 0.119 | 1.372 | 0.244 | 2.963 | 0.089 |
| Weeds, C | **6.888** | **0.010** | **4.249** | **0.042** | 0.127 | 0.722 |
| Sown grasses, C | **8.630** | **0.004** | 3.626 | 0.060 | **13.174** | **0.001** |
| Jaccard similarity | 0.717 | 0.399 | 1.263 | 0.264 | 0.019 | 0.890 |
| **Seed bank** |  |  |  |  |  |  |
| Species richness | **7.321** | **0.008** | **10.186** | **0.002** | 3.606 | 0.061 |
| Total SD | **19.889** | **0.001** | 0.845 | 0.360 | **47.503** | **0.001** |
| Weeds, SD | **62.017** | **0.001** | 0.149 | 0.700 | **76.923** | **0.001** |
| Sown grasses, SD | **17.783** | **0.001** | **4.944** | **0.029** | 2.612 | 0.110 |
| Target species, SD | **42.560** | **0.001** | 2.190 | 0.142 | **43.833** | **0.001** |

**Figure 1.**
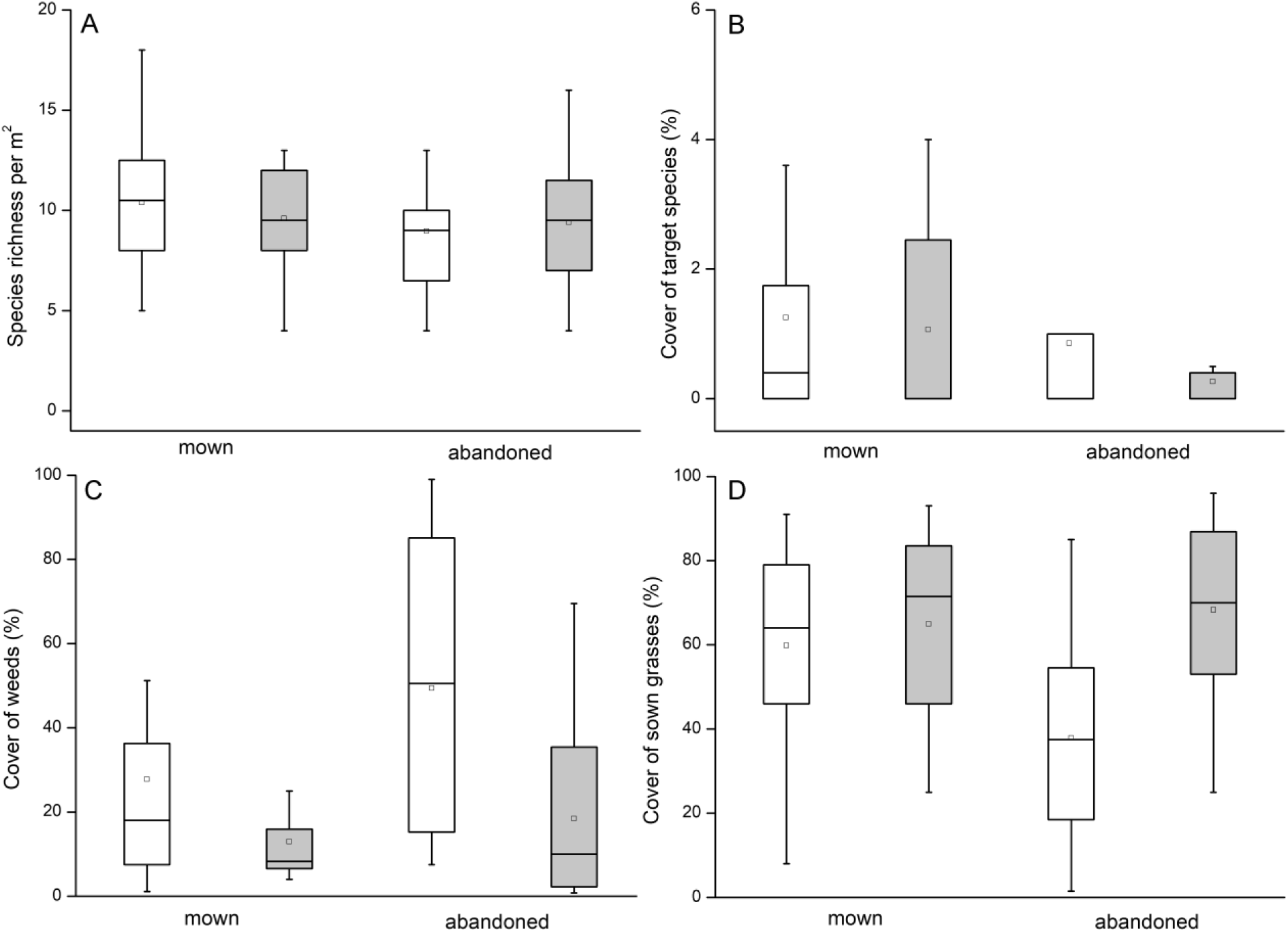
Species richness (A), cover of target species (B), weeds (C), and sown grasses (D) in the vegetation of 8-year-old mown and abandoned restored grasslands. White boxes – restored alkaline grasslands; grey boxes – restored loess grasslands. The boxes show the interquartile range, the lower whiskers show the minimum, the upper whiskers show the maximum, and the inner lines display the median values.

### Seed bank characteristics

The species richness of the seed bank was affected by management and grassland type (Table 1). The highest species richness was recorded in the mown grasslands. Seed density was affected by management and the interaction of management and grassland type (Table 1), being the highest in the mown and alkaline grasslands (Figure 2c). Seed density decreased due to abandonment in the loess grasslands (Figure 2a). In total 5045 seedlings germinated from the seed bank samples. Total seed density ranged between 3183 and 89,127 seeds/m^2^, mean seed density was 13,939 seeds/m^2^. Management and the interaction of management and grassland type affected the seed density of weeds (Table 1). The highest number of weed seedlings was found in the abandoned alkaline grasslands (Figure 2d). Both management and grassland type affected the seed density of sown grasses (Table 1), the highest scores were found in the mown and alkaline grasslands (Figure 2e). The seed density of target species was higher in the mown sites and decreased significantly due to abandonment in the loess grasslands (Table 1, Figure 2f).

**Figure 2.**
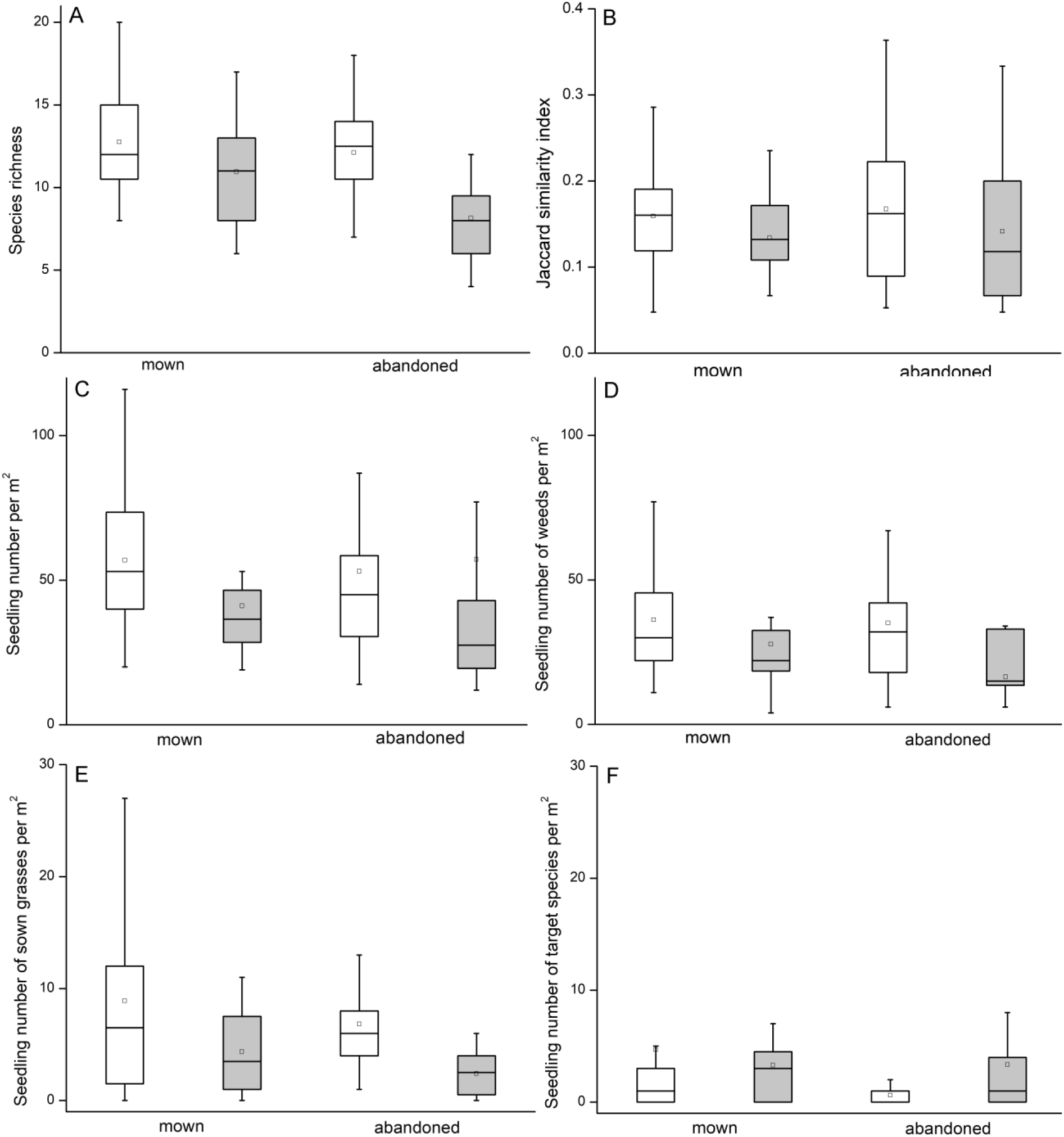
Species richness in the seed bank (A), Jaccard similarity of the vegetation and the seed bank (B), total seedling number (C), the seedling number of weeds (D), sown grasses (E) and target species (F) in the 8-year-old mown and abandoned restored grasslands. Note that one seedling corresponds to a seed density of 265 seeds/m^2^. White boxes – restored alkaline grasslands; grey boxes – restored loess grasslands. The boxes show the interquartile range, the lower whiskers show the minimum, the upper whiskers show the maximum, and the inner lines display the median values.

### Species composition of the vegetation and the seed bank

We found 165 species in the study sites. 106 species were recorded in the vegetation and 129 in the seed bank. 36 species were present only in the vegetation, 59 only in the seed bank and 70 both in the vegetation and the seed bank. The Jaccard similarity of the species composition of the vegetation and seed bank was similarly low (0.16 ± 0.07, mean ± SD) regardless of management and grassland type (Table 1, Figure 2b).

Species composition of the vegetation and the seed bank was clearly separated on the DCA ordination (Figure 3). Seed bank had more homogeneous species composition than the vegetation. In the vegetation, plots sown with alkaline and loess seed mixture were clearly separated. In the seed bank, their separation was not so contrasted. In the vegetation, the species composition of the abandoned plots was more heterogeneous compared to the mown ones; in the seed bank, there was no such trend.

**Figure 3.**
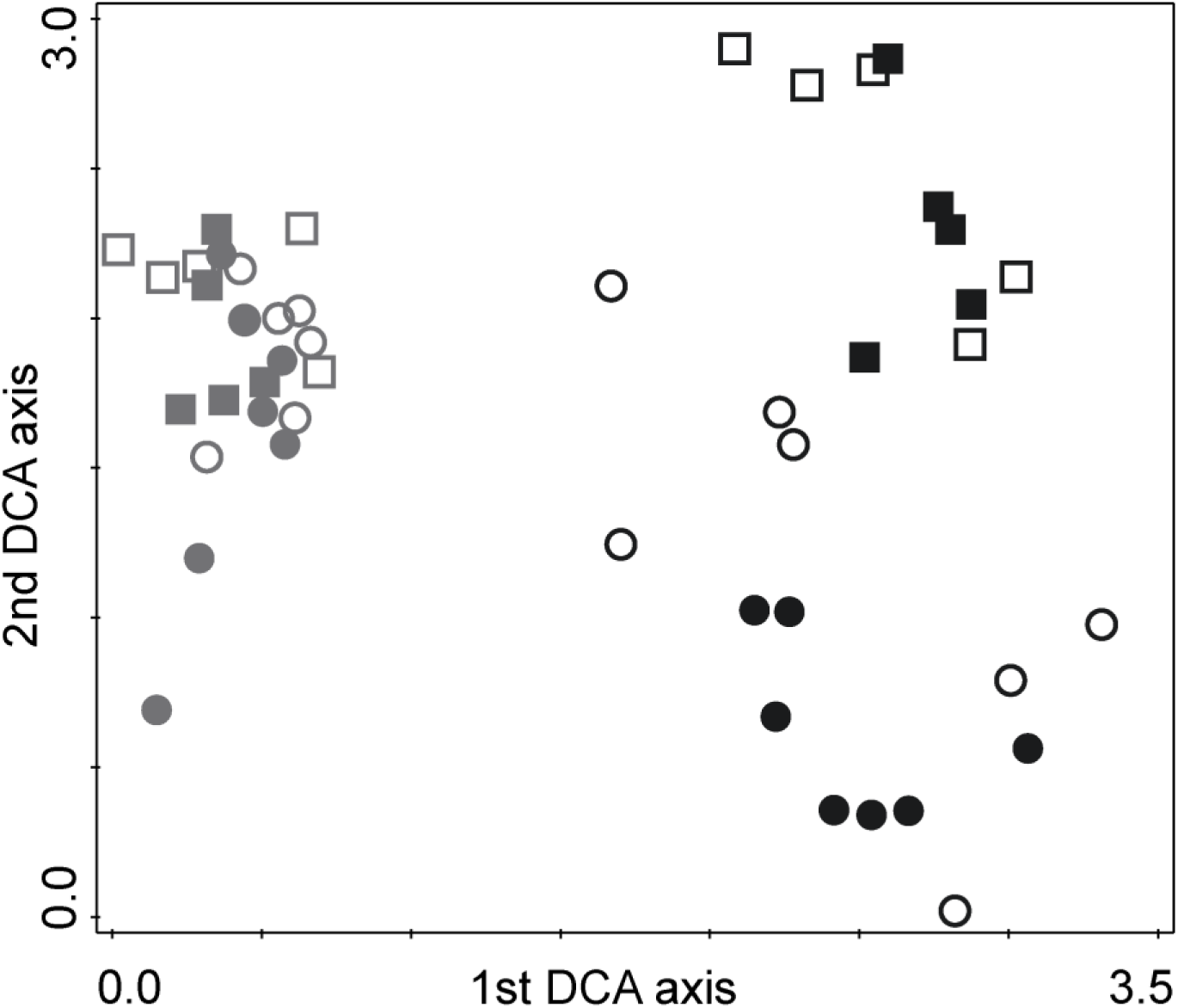
DCA ordination based on the relative abundances of the plant species in the vegetation and seed bank of the restored grasslands. Notations: grey symbols – seed bank, black symbols – vegetation; circles – alkaline grasslands, squares – loess grassland; full symbols – mown sites, empty symbols – abandoned sites. Eigenvalues were 0.5648 (1^st^ axis) and 0.3295 (2^nd^ axis). Cumulative percentage variance of species data was 9.80% for the 1st, and 15.51% for the 2nd axis, respectively.

Both the DCA and the IndVal analysis confirmed that the sown grasses were significant characteristic species to the vegetation of the mown grasslands (*Festuca pseudovina*, and *Poa angustifolia* in the alkaline and *Bromus inermis* and *F. rupicola* in the loess grasslands; Figure 3, Table S1). The mown grasslands’ vegetation included both weeds (*Convolvulus arvensis, Vicia villosa*) and target grassland species (*Achillea collina, Cruciata pedemontana, Trifolium campestre*). There were seven weed species that were characteristic to the vegetation of abandoned grasslands, including *Cirsium arvense, Galium spurium* and *Lactuca serriola*. Characteristic species of the seed bank predominantly included weeds (*Capsella bursa-pastoris, Chenopodium album, Echinochloa crus-gallii*), and a few target species (*Inula britannica, Matricaria chamomilla, Spergularia rubra*; Table S2).

## Discussion

Our study confirmed that the cessation of post-restoration management represents a major threat for the restored grasslands. We found that abandoned grasslands are characterised by a lower cover of perennial grasses and higher cover of weeds compared to mown ones, which partly confirmed our first hypothesis. We did not detect a decline in species richness due to abandonment, which is likely due to the generally low species richness of the studied mown and abandoned grasslands. The most abundant weed species of the abandoned grasslands was *Cirsium arvense*, which is among the most dangerous weed species worldwide and the third most noxious weed in Europe (Friedli & Bacher 2001). Its high cover in the vegetation of abandoned grasslands represents a major threat of future encroachment, as the species is known to produce a large amount of long-term persistent seeds (Tiley 2010).

We found that abandoned alkaline and loess grasslands showed distinct vegetation composition: alkaline grasslands were mainly characterized by weeds while loess grasslands were characterised by the high cover of the sown grasses. This is partly confirmed our second hypothesis as we found that grassland type affected the vegetation and seed bank of abandoned grasslands. We expected that restored loess grasslands are more sensitive to the effects of abandonment, because contrary to alkaline grassland species, most species characteristic to natural loess grasslands do not have persistent seed bank (Tóth & Hüse 2014, Valkó et al. 2014). Even though we detected higher seed density in the alkaline grasslands, the expression of the seed bank was likely hampered by the strong biotic filtering effect of the sown grasses (Deák et al. 2011). Also, natural loess grasslands are generally very sensitive to the changes in their management regimes (Kelemen et al. 2013). However, in our study the cover of perennial grasses remained high in abandoned loess grasslands. This can be attributed to the biotic filtering effect of one of the sown grass species, *Bromus inermis*, which is a highly competitive species that could persist in the loess grasslands regardless of abandonment and could effectively suppress weeds in the vegetation (see also Kelemen et al. 2014). However, on the long run, the encroachment of *B. inermis* can lead to the decreased diversity of the loess grasslands as it might competitively exclude other target species from the vegetation.

Our study confirmed the limited potential of soil seed bank in the maintenance of species richness of the restored alkaline and loess grasslands. In general, the seed bank was dominated by weeds and there were only a few target species (see also Klaus et al. 2018; Wagner et al. 2018). The similarity of the species composition of vegetation and seed bank was low, as was found in other restored grasslands (Rayburn et al. 2016; Godefroid et al. 2018). Thus, it is likely that abandonment affects the vegetation and seed bank through different mechanisms. Contrary to our third hypothesis, we found that five years of abandonment had a larger effect on the seed bank than on the vegetation. Abandonment had a significant effect on all seed bank characteristics. In abandoned grasslands, we found a decreased species richness and seed density, increased seed density of weeds and lower seed density of sown grasses and target species. In general, the seed bank of the restored grasslands was characterised by annual weeds (more than 70% of all viable seeds), such as *Capsella bursa-pastoris, Chenopodium album* and *Conyza canadensis*. The seed bank of weeds followed the vegetation changes after abandonment (see also Shang et al. 2016); seed density of weeds increased in the alkaline and decreased in the loess grasslands. This might be connected to the high cover of *Bromus inermis* in the loess grasslands, which probably prevented the establishment and seed production of weed species and therefore decreased the rate of the build-up of their seed bank. Contrary, in abandoned alkaline grasslands, the cover of weeds increased in the vegetation, which contributed to the build-up of their seed bank. This was supported by the decreased cover of target grass species in the abandoned sites.

Our findings demonstrated that post-restoration management is important to maintain the cover of sown grasses and suppress weed species. We found that abandonment leads not only to the encroachment of weed species in the vegetation, but also to the build-up of their seed bank; these two synergic processes pose a considerable threat for the sustainability and ecosystem functioning of restored grasslands. In regions where animal husbandry is economically profitable, the long-term management of the restored meadows and pastures can be ensured (Abson et al. 2017). In other cases, the management of restored grasslands is often guaranteed only in the short term, which is generally limited to the few-year-long maintenance period of the restoration project (Valkó et al. 2018). Our results highlight that it is inevitable to design the long-term management of the sites subjected to grassland restoration already in the planning phase of the restoration projects and ensure that the management plan is ecologically and economically feasible.

## Acknowledgements

We are thankful to the colleagues of Hortobágy National Park Directorate (L. Gál, I. Kapocsi) for their help in fieldwork. Authors were supported by NKFI KH 126476 (OV), NKFI FK 124404 (OV), NKFI KH 130338 (BD), OTKA K 116239 (BT), NKFI KH 126477 (BT), NKFIH K 119225 (PT), NKFI PD 124548 (TM), NKFI PD 128302 (KT), MTA’s Post Doctoral Research Program (AK) and the Bolyai János Fellowship of the Hungarian Academy of Sciences (OV, BD, AK).

